# Transgenerational plasticity of inducible defenses: combined effects of grand-parental, parental and current environments

**DOI:** 10.1101/589945

**Authors:** Juliette Tariel, Sandrine Plénet, Émilien Luquet

## Abstract

While an increasing number of studies highlights that parental environment shapes offspring phenotype (transgenerational plasticity TGP), TGP beyond the parental generation has received less attention. Studies suggest that TGP impacts population dynamics and evolution of phenotype, but these impacts will depend on how long an environmental effect can persist across generations and whether multigenerational effects are cumulative. Here we tested the impact of both grand-parental and parental environments on offspring reaction norm in a prey-predator system. We exposed three generations of *Physa acuta* snails without and with predator-cues according to a full factorial design and measured offspring inducible defenses. We found that both grand-parental and parental exposure to predator cues impacted offspring anti-predator defenses, but their effects were not cumulative and depended on the defenses considered. We also highlighted that grand-parental environment could alter reaction norm of offspring shell thickness, demonstrating an interaction between the grand-parental TGP and the within-generational WGP plasticity. We called for more studies covering the combine effects of multigenerational environments.

## Introduction

Organisms may respond to fluctuating environments by adapting through genetic evolution over generations or through phenotypic plasticity. This last is traditionally defined as the capacity of a given genotype to render alternative phenotypes under different environmental conditions (within-generational plasticity, WGP) (West-Eberhard 2003; Pigliucci 2005). Plasticity may also occur across generations (transgenerational plasticity, TGP), when the phenotype of offspring is influenced by carry-over effects of past environments experienced by the previous generation(s) (Agrawal et al. 1999; Galloway and Etterson 2007; Salinas et al. 2013). Ancestors can alter the phenotype of their offspring without involving genetic changes through a range of non-genetic processes as parental effects, *e*.*g*. transmission of nutrients, hormones, proteins (Mousseau 1998; Crean and Bonduriansky 2014), or by any form of epigenetic inheritance, *e*.*g*. DNA methylation marks, histone protein modifications, non-coding small RNAs (Holeski et al. 2012; Schlichting and Wund 2014). TGP has been shown for diverse organisms and traits in response to various environments (Bonduriansky and Day 2009; Salinas et al. 2013; Donelson et al. 2018). When past environmental cues provide reliable proxies about future offspring conditions, TGP may enable organisms to cope with fast-changing environments because it refines offspring phenotypes in anticipation of the environmental conditions they are likely to experience (Bonduriansky and Day 2009; Herman and Sultan 2011; Donelson et al. 2018).

Most studies on TGP focused on the effect of parental environment on offspring phenotype (*e*.*g*. Mousseau 1998; Wolf and Wade 2009; Herman and Sultan 2011; Donelson et al. 2018) and on offspring reaction norm (interaction TGP x WGP; Salinas et al. 2013). But transgenerational effects are not confined to the parental generation. A few examples have shown that TGP can persist for multiple successive generations (Plaistow et al. 2006; Remy 2010; Sarker and Peleg-Raibstein 2019), even though the majority is restrained to three generations (*i*.*e*. grand-parental effect) (*e*.*g*. Hafer et al. 2011; Herman and Sultan 2011; Kou et al. 2011; Lock 2012; Shama et al. 2014; Walsh et al. 2014). However, it is unclear how multigenerational effects can interact (Prizak et al. 2014) and experimental works controlling for combination of multigenerational effects always showed complex patterns of phenotypic offspring responses. For example, Hafer et al. (2011) and Walsh et al. (2014) in collembolan (*Folsomia candida*) and cladoceran (*Daphnia ambigua*) respectively, demonstrated that some life-history traits were affected by an interactive effect between grandparental and parental environment, leading to phenotypic landscapes that do not fit simple adaptive scenarios. Moreover, such complex patterns of phenotypic responses can depend on the offspring context. Plaistow et al. (2006), for instance, showed in the soil mite *Sancassania berlesei* that the persistence of past environments (across four generations) differed between high- and low-food offspring contexts. Consequently, understanding how multigenerational combined environments interact to shape the offspring phenotype is still limited and challenging.

Predator-induced plasticity is a well-known model in WGP study (*e*.*g*. Harvell 1990; Relyea 2001; Hoverman et al. 2005) and allows an individual to fine-tune its phenotype facing predation risk (Lima 1998; Tollrian and Harvell 1999; Benard 2004). Predator-induced defenses are also widely used to investigate TGP over two generations (parental and offspring generations) (*e*.*g*. Agrawal et al. 1999; Bell and Stein 2017; Colicchio 2017). However, to our knowledge, only two studies have been interested in TGP of predator-induced traits beyond two consecutives generations (Walsh et al. 2015; Sentis et al. 2018).

We focused on a hermaphroditic gastropod *Physa acuta*. Physidae are well-known to develop adaptive phenotypes in response to predation risk (DeWitt et al. 1999; Auld and Relyea 2008, 2011; Gustafson et al. 2014; Auld and Houser 2015; Beaty et al. 2016). Predator (crayfish) cues induce WGP of *Physa sp*. life-history traits (larger age and size at first reproduction; Auld and Relyea 2008), shell thickness (thicker shell; Auld and Relyea 2011), shell size (narrower shape; DeWitt 1998), and behavior (crawling-out the water; Alexander and Covich 1991; DeWitt et al. 1999). We exposed three successive generations of snails from hatching to sexual maturity according to a full factorial design to environments without and with predator-cues. The results concerning the first two generations have demonstrated a predator-induced TGP in *P*. *acuta* (Luquet and Tariel 2016) that has been confirmed in a concomitant study (Beaty et al. 2016). Here, we focused on the F3 generation to investigate how the effects of grand-parental, parental and offspring environments combine to influence behavior, shell thickness and shell morphology.

## Methods

### Animal collection and experimental design

Adult *P. acuta* snails were collected (the experimental design represented on Appendix 1) on March 2015 in a lentic backwater of the Rhône river (45° 48’6”N, 4° 55’33”E) in Lyon, France. *Physa acuta* is a freshwater and simultaneous hermaphroditic snail, invasive from North America (Lydeard et al. 2016). The wild-caught adult snails constituted our F_0_ generation. We pooled them overnight in a 10L-aquarium to ensure that offspring result from outcrossing (*P. acuta* is a preferential outcrosser; Jarne et al. 2010). Then, we individually isolated all F_0_ snails in 70 mL plastic vials filled with reconstituted water (2.4 g NaHCO_3_, 3 g CaSO_4_, 1.5 g MgSO_4_, 0.1 g KCl to 25 L deionized water) in a 25°C experimental room with 12h light-dark photoperiod. After 24 hours, we removed the F_0_ adults from the vials and we randomly choose 15 vials with one egg capsule each. These 15 egg capsules constituted our 15 maternal families (hereafter called only “families”) of the F_1_ generation and developed until hatching (∼ 7 days). Two days after hatching, we randomly sampled 12 siblings per family and split them into two environments: 6 snails remained in a no-predator environment (control environment) while 6 others were moved in a predator-cue environment. These F_1_ snails were reared in 70 mL plastic vials with their siblings until 28 days-old where they were isolated in the same type of plastic vials until the end of the experiment (35 days after hatching). We then generated the second F_2_ generation in merging F_1_ snails into copulation groups of 15 individuals (one snail from each of the 15 families). At this step, the snail identity was not recorded and thus the pedigree could not be further assessed. We made 6 copulation groups per treatment and each reproduced in a 5L-aquarium for 24h in no-predator water to ensure embryos are not exposed to predator environment. Then, individuals were isolated, and we randomly selected 18 F_1_ snails that had laid eggs from each treatment. We then followed the same protocol as previously to rear F_2_ snails in control and predator-cue environments according to a full factorial design until 49 days after hatching. The F_3_ generation was then generated and reared using the procedure described above. As growth rate was slowing down every generation under our laboratory conditions and we wanted snails in a reasonably sufficient size, we reared the F_3_ generation at a later age (74 days after hatching). This F_3_ generation was represented by eight combinations of grand-parental (E_1_), parental (E_2_) and offspring (E_3_) environments: CCC, CPC, PCC, PPC, CCP, CPP, PCP and PPP with each time “C” for control environment and “P” for predator-cue environment (Figure 1). The number of individuals and families per combination of environments are reported on Figure 1.

**Figure 1.**
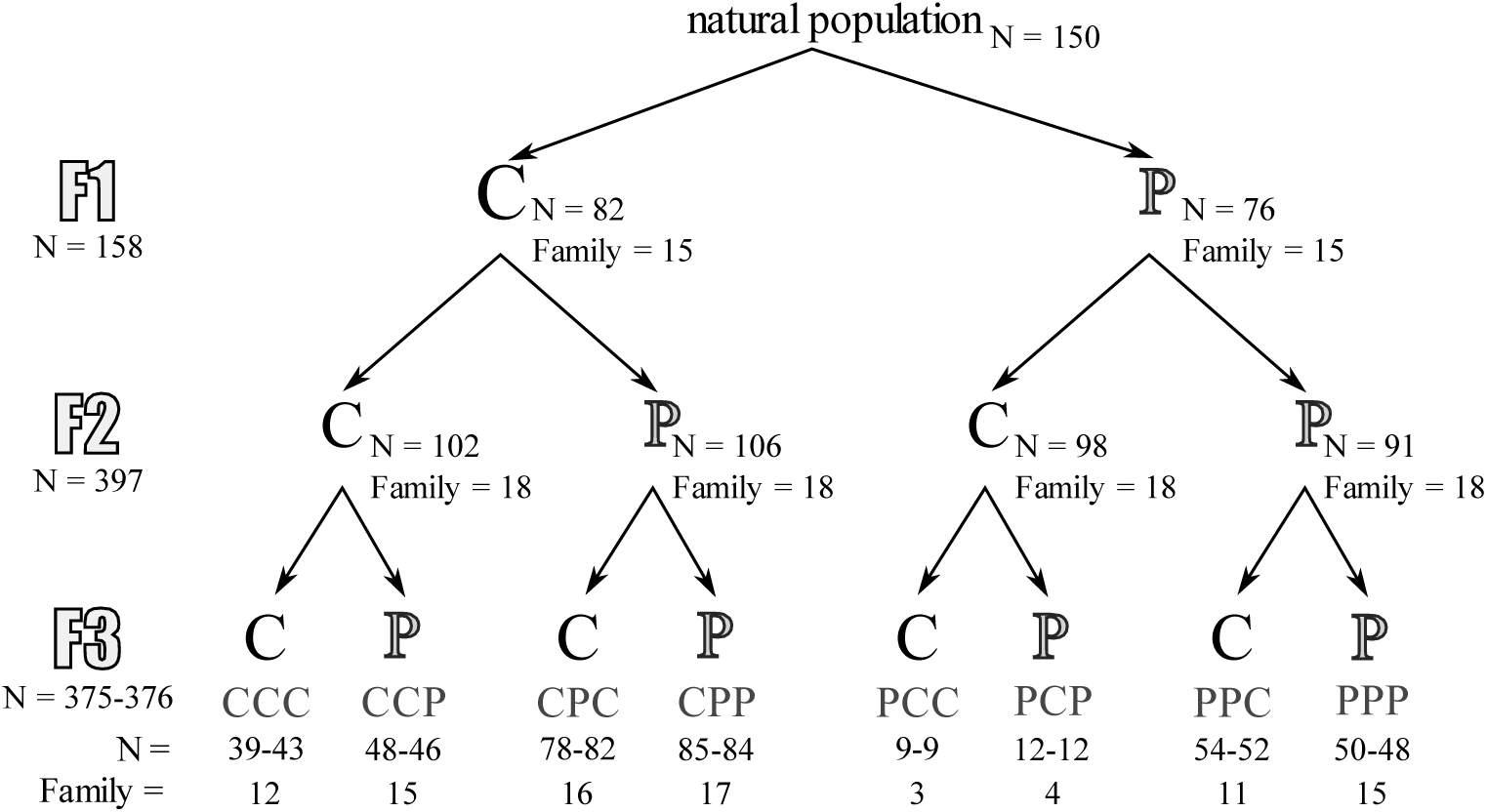
Number of individuals (N) and families (Family) at each generation and in the two treatments with “C” for control environment and “P” for predator-cue environment. For the 3^rd^ generation, two number of individuals are reported, one for behavioral measurements (first position) and one for other measurements (second position).

Water and food (*ad libitum*, chopped and boiled lettuce) were renewed for all experimental snails twice a week. Predator-conditioned water with predator cues was obtained by individually rearing crayfishes (*Procambarus clarkii*) in 4L reconstituted water and feeding with one *P. acuta* adult one day before water change (Auld and Relyea 2011). This crayfish-conditioned water was used for the predator-cue treatment while only reconstituted water was used for the control treatment. This crayfish species coexists with *P. acuta* in its native location in North America.

### Measuring phenotypes

We assessed anti-predator behavior three times in 70-day-old F_3_ snails through three consecutive days starting one day after the water change. We measured anti-predator behavior by recording the position above/on or below the water surface in the rearing boxes with predator cues present or absent according to the treatment. Crawling-out of the water (position above water surface) is considered as allowing to escape from benthic predators like crayfishes (DeWitt et al. 1999).

At 74 days old, we gently dried snails with paper towel and measured the snail total wet weight (body and shell) with an electronic scale at the nearest 0.001mg. A photograph of each snail aperture upwards was taken with an Olympus SC50 camera installed on an Olympus SZX9 binocular and its Olympus DF PLAPO 1X-2 objective at a x8 magnification. Shell and aperture length and width (shell morphology) were measured on these photographs with the software ImageJ (Schneider et al. 2012). Shell thickness was measured with an electronic caliper at the nearest 0.01mm at the edge of the aperture. Shorter and narrower shell and aperture dimensions (after adjusting for mass) and thicker shell are adaptive anti-predator responses (Auld and Relyea 2011).

### Statistical analysis

The multigenerational effect of predator cues on anti-predator behavior (*i*.*e*. snail position above/on or below the water surface) was analyzed using generalized linear mixed models (GLMM) assuming a binomial distribution (logit link function). Shell and aperture length and width were analyzed with a principal component analysis (PCA) with the package FactoMineR (Lê et al. 2008) to assess shell morphology. They were highly correlated and the first component (PC1) explained most of variance in shell morphology (95.8%; Appendix 2): a high value of PC1 was associated with a longer and wider shell and aperture, whereas a low value was associated with a smaller and narrower shell and aperture. PC1 was then used as a proxy for shell morphology and response variable in linear mixed model. The multigenerational effect of predator cues on weight, shell thickness and shell morphology was analyzed using linear mixed models (LMM) with restricted maximum likelihood estimation and Kenward and Roger’s approximation for degrees of freedom. These response variables were scaled prior to statistical analysis. In all linear mixed models, grand-parental (E_1_), parental (E_2_), offspring (E_3_) environments, and all interactions were considered as fixed effects. Because shell thickness and shell morphology were significantly correlated with weight, weight (cube root transformed and then scaled) was added as a continuous fixed effect (analyses of covariance) in these models. Family was considered as a random intercept effect in all models. Individual identity was added as a random intercept model in the GLMM model (anti-predator behavior analysis) to account for repeated measures on the same individual. We used type 2 F-tests for significance of fixed effects in the LMM and likelihood ratio tests in the GLMM (lmerTest package; Kuznetsova et al. 2017). We tested significance of random effects with likelihood ratio tests comparing models with or without the tested random effect in the full fixed effect structure. All statistical analyses were performed with R 3.4.1 (R Core Team 2017) and the glmer() and lmer() functions from the lme4 package (Bates et al. 2015 p. 4).

## Results

### Anti-predator behavior

The offspring exposure to predator cues (E_3_) significantly increased by 105% the proportion of snails crawling-out the water (Table 1a; Fig. 2a). The parental environment (E_2_) did not affect the proportion of crawling-out behavior (Table 1a; Fig. 2a). However, grand-parental exposure to predator cues (E_1_) significantly increased by 28 % the proportion of crawling-out behavior (Table 1; Fig. 2a). Family and individual random effects were significant (Table 1a).

**Table 1.** Results of generalized mixed model (**a**) and linear mixed model (**b, c** and **d**) analysis on offspring predator-induced traits. Bold values indicate significant P-values (P < 0.05).

|  |  |  |  |  |  |
| --- | --- | --- | --- | --- | --- |
| a. Crawling-out | *Fixed effects* | *Estimate (SE)* | *Df* | *Chisq* | *P* |
| | Grand-parental env. ($E_1$) | 0.848 (0.6812) | 1 | 8.29 | **0.0040** |
| | Parental env. ($E_2$) | -0.090 (0.3864) | 1 | 0.4 | 0.5274 |
| | Offspring env. ($E_3$) | 1.738 (0.3544) | 1 | 114.46 | **<0.0001** |
| | $E_1 \times E_2$ | -0.389 (0.7696) | 1 | 0.01 | 0.9315 |
| | $E_1 \times E_3$ | -0.168 (0.7882) | 1 | 1.89 | 0.1687 |
| | $E_2 \times E_3$ | -0.146 (0.4242) | 1 | 0.01 | 0.9072 |
| | $E_1 \times E_2 \times E_3$ | 0.855 (0.8915) | 1 | 0.91 | 0.3395 |
|  | *Random effects* | *Variance* | *Df* | *Chisq* | *P* |
|  | Family | 0.391 | 1 | 14.98 | **0.0001** |
|  | Individual | 0.489 | 1 | 8.26 | **0.0040** |
| b. Weight | *Fixed effects* | *Estimate (SE)* | *Numdf, Dendf* | *F* | *P* |
| | Grand-parental env. ($E_1$) | -0.047 (0.1656) | 1, 42.44 | 0.18 | 0.6766 |
| | Parental env. ($E_2$) | -0.116 (0.1656) | 1, 48.70 | 0.57 | 0.4545 |
| | Offspring env. ($E_3$) | -0.366 (0.1329) | 1, 354.26 | 14.76 | **0.0001** |
| | $E_1 \times E_2$ | -0.041 (0.3312) | 1, 55.77 | 0.01 | 0.9105 |
| | $E_1 \times E_3$ | -0.027 (0.2659) | 1, 348.77 | 0.001 | 0.9737 |
| | $E_2 \times E_3$ | -0.104 (0.2659) | 1, 362.32 | 0.30 | 0.5836 |
| | $E_1 \times E_2 \times E_3$ | 0.069 (0.5318) | 1, 358.29 | 0.02 | 0.8969 |
|  | *Random effects* | *Variance* | *Df* | *Chisq* | *P* |
|  | Family | 0.085 | 1 | 6.6 | **0.0102** |
| c. Shell thickness | *Fixed effects* | *Estimate (SE)* | *Numdf, Dendf* | *F* | *P* |
|  | cuberoot(Weight) | 0.656 (0.0404) | 1, 364.6 | 257.77 | **<0.0001** |
| | Grand-parental env. ($E_1$) | -0.092 (0.1152) | 1, 40.41 | 0.19 | 0.6650 |
| | Parental env. ($E_2$) | 0.057 (0.1153) | 1, 49.59 | 0.04 | 0.8446 |
| | Offspring env. ($E_3$) | 0.258 (0.1054) | 1, 356.90 | 28.15 | **<0.0001** |
| | $E_1 \times E_2$ | 0.180 (0.2305) | 1, 59.10 | 0.62 | 0.4329 |
| | $E_1 \times E_3$ | -0.350 (0.2088) | 1, 350.26 | 4.01 | **0.0461** |
| | $E_2 \times E_3$ | 0.497 (0.2088) | 1, 363.53 | 7.68 | **0.0059** |
| | $E_1 \times E_2 \times E_3$ | 0.033 (0.4176) | 1, 359.10 | 0.01 | 0.9366 |
|  | *Random effects* | *Variance* | *Df* | *Chisq* | *P* |
|  | Family | 0.020 | 1 | 0.78 | 0.3780 |
| d. Shell morphology | *Fixed effects* | *Estimate (SE)* | *Numdf, Dendf* | *F* | *P* |
|  | cuberoot(Weight) | 1.901 (0.0167) | 1, 366.61 | 12842.53 | **<0.0001** |
| | Grand-parental env. ($E_1$) | -0.034 (0.0496) | 1, 41.10 | 0.19 | 0.6613 |
| | Parental env. ($E_2$) | -0.014 (0.0497) | 1, 49.12 | 0.33 | 0.5677 |
| | Offspring env. ($E_3$) | -0.287 (0.0433) | 1, 355.56 | 73.25 | **<0.0001** |
| | $E_1 \times E_2$ | -0.044 (0.0992) | 1, 57.48 | 0.12 | 0.7288 |
| | $E_1 \times E_3$ | 0.053 (0.0858) | 1, 349.39 | 0.02 | 0.8943 |
| | $E_2 \times E_3$ | 0.080 (0.0859) | 1, 362.82 | 3.68 | 0.0558 |
| | $E_1 \times E_2 \times E_3$ | -0.215 (0.1717) | 1, 358.51 | 1.57 | 0.2114 |
|  | *Random effects* | *Variance* | *Df* | *Chisq* | *P* |
|  | Family | 0.005 | 1 | 1.88 | 0.1704 |

**Figure 2.**
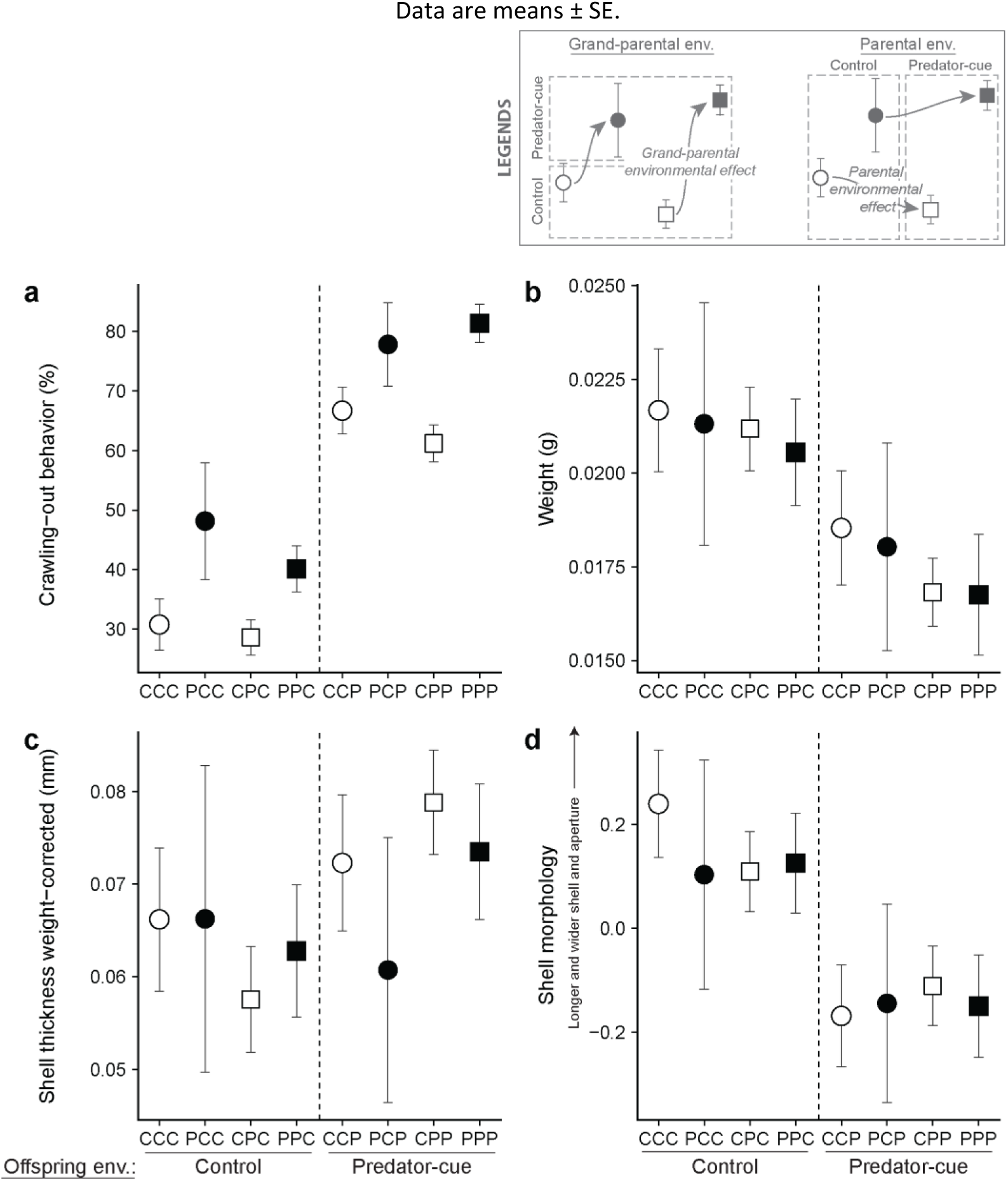
The effect of multi-generational exposure to predator-cues on offspring **a** crawling-out behavior (proportion of snails out the water in %), **b** total weight, **c** and **d** least-mean squares of shell thickness and shell morphology in the model accounting for weight described in Table 1. CCC, PCC, CPC, PCC, CCP, PCP, CPP and PPP represent the eight combinations of grand-parental (E_1_), parental (E_2_) and offspring (E_3_) environments with “C” for control environment and “P” for predator-cue environment for every generation. The vertical dashed line separates the two offspring treatment groups. White shapes are for grand-parental control environment and black shapes for grand-parental predator-cue environment. Circles are for parental control environment and squares for parental predator-cue environment.

### Snail weight

The offspring exposure to predator cues significantly decreased by 18% the snail weight (Table 1b; Fig. 2b). Neither the parental environment nor the grand-parental environment influenced snail weight (Table 1b; Fig. 2b). Family random effect was significant (Table 1b).

### Shell thickness

After accounting for snail weight, grand-parental and parental environments interacted both with the offspring environment (Table 1c; Fig. 2c), demonstrating that both could affect the offspring response to predator cues. In the offspring control environment, the grand-parental exposure to predator cues increased by 4% the offspring shell thickness whereas the parental exposure to predator cues decreased by 9% the offspring shell thickness. And in the offspring predator-cue environment, the grand-parental exposure to predator cues decreased by 11% the offspring shell thickness whereas the parental exposure to predator cues increased by 14% the offspring shell thickness. Regarding the direct effect of offspring environment, offspring from predator-cue environment had a thicker shell than those from current control environment (Table 1c; Fig. 2c). Family random effect was not significant (Table 1c).

### Shell morphology

After accounting for snail weight, the offspring exposure to predator cues impacted the shell morphology and induced a shorter and narrower shell and aperture (Table 1d; Fig. 2d). Neither the parental environment (interaction parental and offspring environments almost significant P < 0.10) nor the grand-parental environment influenced the shell morphology (Table 1d; Fig. 2d). Family random effect was not significant (Table 1d).

## Discussion

We first confirm that the exposure to predator cues induces well-known defenses against crayfish predation in *P. acuta*. Our key finding is that predator cues alter also offspring defenses two generations later but depending on the offspring environment (WGP x TGP) and the defensive traits considered. Our experimental work highlights that TGP effects can be complex beyond the parental generation and that the offspring phenotype results from a combination of multigenerational effects.

Corroborating several studies on anti-predator defenses of *Physa sp*. (DeWitt et al. 1999, 2000; Turner et al. 1999; Dalesman et al. 2009; Auld and Relyea 2011), the offspring exposure to predator cues induced higher crawling-out behavior, shell-crushing resistance (thicker shell) and entry-resistant shell (narrower shell and aperture). Moreover, offspring exposed to predator cues were lighter, suggesting a trade-off, *i*.*e*. a lower energetic investment in growth due to a potential cost to produce these defenses (as showed in other gastropod species: Brönmark et al. 2012). This result stresses the fitness advantage of WGP which allow the production of costly defenses only in case of predation (Harvell 1990).

### Grand-parental and parental effects on anti-predator defenses

TGP is expected to evolve when the ancestral environment is a good proxy of offspring environment (Harvell 1990; Uller 2008; Bonduriansky and Day 2009; English et al. 2015; Leimar and McNamara 2015; Dey et al. 2016), allowing a pre-adaptation of offspring to predation risk (Agrawal et al. 1999). In our predator-prey system, crayfish has a long lifespan (*ca*. 3 years) compared to the generation time of *P. acuta* (*ca*. 50 days) and a relatively sedentary lifestyle (Vioque-Fernández et al. 2009). This suggests that generational cues of crayfish presence can be a good proxy of predation risk across several snail generations and thus that TGP could have long-lasting effects on the anti-predator defenses. Consistently, in *P. acuta*, parental exposure to predator cues induces a more crush-resistant shell and an higher escape behavior in offspring (Beaty et al. 2016; Luquet and Tariel 2016). In this study, as expected, TGP went further than the parental generation: the grand-parental environment also influenced the anti-predator behavior and the shell thickness of offspring.

How long can persist transgenerational effects on anti-predator responses is a remaining open question. To our knowledge, the study of Sentis et al. (2018) on the pea aphid (*Acyrthosiphon pisum*) is the only one to investigate predator-induced TGP over a large number of generations (> 25). They found that the defensive phenotype – a high frequency of winged aphids in the population – persists for one generation after removing predators whatever the induction time is, *i*.*e*. the previous number of successive generations experiencing the novel environment (predator presence). However, three generations are needed after removing predators for the frequency of winged phenotypes to come back to the control level, and this number of generations increases with the induction time. Together, these results suggest that multigenerational environmental effects on inducible defenses are broader than just a parental effect and could persist for many generations.

### Combination of multigenerational effects on anti-predator defenses

We showed that the offspring phenotype results from a combination of multigenerational effects (grand-parents, parents and offspring), similar to theoretical and other experimental studies (Hafer et al. 2011; Kou et al. 2011; Lock 2012; Prizak et al. 2014; Shama and Wegner 2014; Walsh et al. 2014). However, in our study, grand-parental and parental effects acted independently (no significant interaction between grand-parental and parental environmental effects): either only one affected the offspring environment (behavior), or in interaction with the offspring environment (WGP x TGP) and in opposite directions (shell thickness). This results in complex offspring phenotypic patterns that do not fit with simple anti-predator scenarios. The adaptive relevance of such combinations of multigenerational effects is far to be evident even studying, as in our study, fitness-related traits. It would be thus interesting to compare the survival of snails from different past environmental histories exposed to lethal predation challenges. The offspring crawling-out behavior increased with offspring and grand-parental exposure to predator cues while the parental environment did not alter this behavior. Shell thickness was influenced by both grand-parental and parental environments, but in opposing directions and depending on the offspring environment (grand-parental and parental WGP x TGP interactions). In offspring control environment, grand-parental exposure to predator cues increased the offspring shell thickness whereas parental exposure reduced the shell thickness. The effects were opposite in the offspring predator-cue environment.

Firstly, these results confirm that offspring reaction norms can be altered by parental environment (shell thickness; Salinas et al. 2013; Luquet and Tariel 2016; Donelson et al. 2018) but expand for the first time the WGP x TGP interaction to grand-parental environmental cues. Secondly, opposing directions of grand-parental and parental effects found on shell thickness are not rare in empirical studies (Magiafoglou and Hoffmann 2003; Shama and Wegner 2014) and illustrate the complexity to determine the adaptive significance of multigenerational effects. Such opposing effects may reflect different mechanisms underlying the transfer of environmental information (Shea et al. 2011). This complex opposing relationship between grand-parental and parental environmental effects could be also theoretically beneficial by reducing the phenotypic variance which allow the population to stay closer to the target phenotype (Prizak et al. 2014). Moreover, focusing on few generations in short-term experiments artificially focuses the interpretations of such effects while they could only be transient over larger timescales in a population dynamic framework. For example, Sentis et al. (2018), after removing predators, observed that the frequency of winged aphids remained high for one generation before dropping abruptly below the control levels (grand-parental effect), and then converging with the winged aphid frequencies of the control lines (great-grand-parental effect). Consequently, in focusing on only three consecutive generations as in our study, these results could be interpreted as a negative grand-parental effect (decrease of winged aphid frequency) opposing to a positive parental effect (increase of winged individual frequency) on the offspring phenotype. These findings highlight the need to develop empirical studies on larger timescales and controlling for the combination of multigenerational effects.

### Trait-dependence of transgenerational plasticity

Our results show that the pattern of TGP depends on the traits (anti-predator behavior, shell thickness and shell morphology). Behavioral traits, which are often labile and exhibiting reversible WGP within developmental or adult stages, are predicted to be influenced by current environment rather than by past environmental experience (Piersma and Drent 2003; Dingemanse and Wolf 2013). Behavioral WGP in response to current environmental cues should rapidly by-pass the behavioral TGP (Beaman et al. 2016). By contrast, the traits that are more constrained during the development and exhibiting irreversible variations, as morphological traits, are predicted to be relatively more influenced by past environments (Kuijper and Hoyle 2015). TGP on morphological traits could irreversibly engage the offspring on developmental trajectories and could not be compensated by WGP. In *P. acuta*, crawling-out behavior is indeed very flexible and reversible at a time scale of hours while a thicker shell and a narrower shell shape are irreversible changes in the developmental trajectory (DeWitt et al. 1999; Relyea 2003). Surprisingly in our study, the escape behavior of offspring is influenced by the grand-parental environment while shell morphology was not influenced by parental or grand-parental environments. This highlights that transgenerational effects on morphological traits may have a short persistence over generations while behavioral TGP may be much more prevalent than currently realized. Parental TGP on behavioral traits has been sometimes observed (*e*.*g*. Storm and Lima 2010; Giesing et al. 2011; Bestion et al. 2014; Donelan and Trussell 2015) and few times with long-lasting effects over generations (Dias and Ressler, 2014; Remy, 2010).

### Conclusion

In our study, we demonstrated that the effects of multigenerational (grand-parental, parental and offspring) exposure to predator cues on a variety of offspring defensive traits (escape behavior, shell thickness and shape) do not fit simple anti-predator scenarios. The multigenerational effects combined, sometimes in opposing directions and depending on the traits, to shape the offspring anti-predator defenses. We also call for more theoretical and empirical studies integrating the combined effects of multigenerational environments on larger number of generations, investigating the underlying mechanisms (epigenetic inheritance) and evaluating their evolutionary importance.

## Data and code accessibility

Data and R script are available from the Zenodo repository (https://doi.org/10.5281/zenodo.2687257).

## Acknowledgements

Thanks to P. Gibert for advice on the manuscript. We thank L. Bensousoussan for help on data analysis. We are grateful to the PCI recommender Troy Day as well as two reviewers (Stewart Plaistow and an anonymous reviewer) for their useful and high-quality reviews based on a previous version of the manuscript. This preprint has been reviewed and recommended by Peer Community In Evolutionary Biology (https://doi.org/10.24072/pci.evolbiol.100076).

## Appendix

### Appendix 1. Experimental design

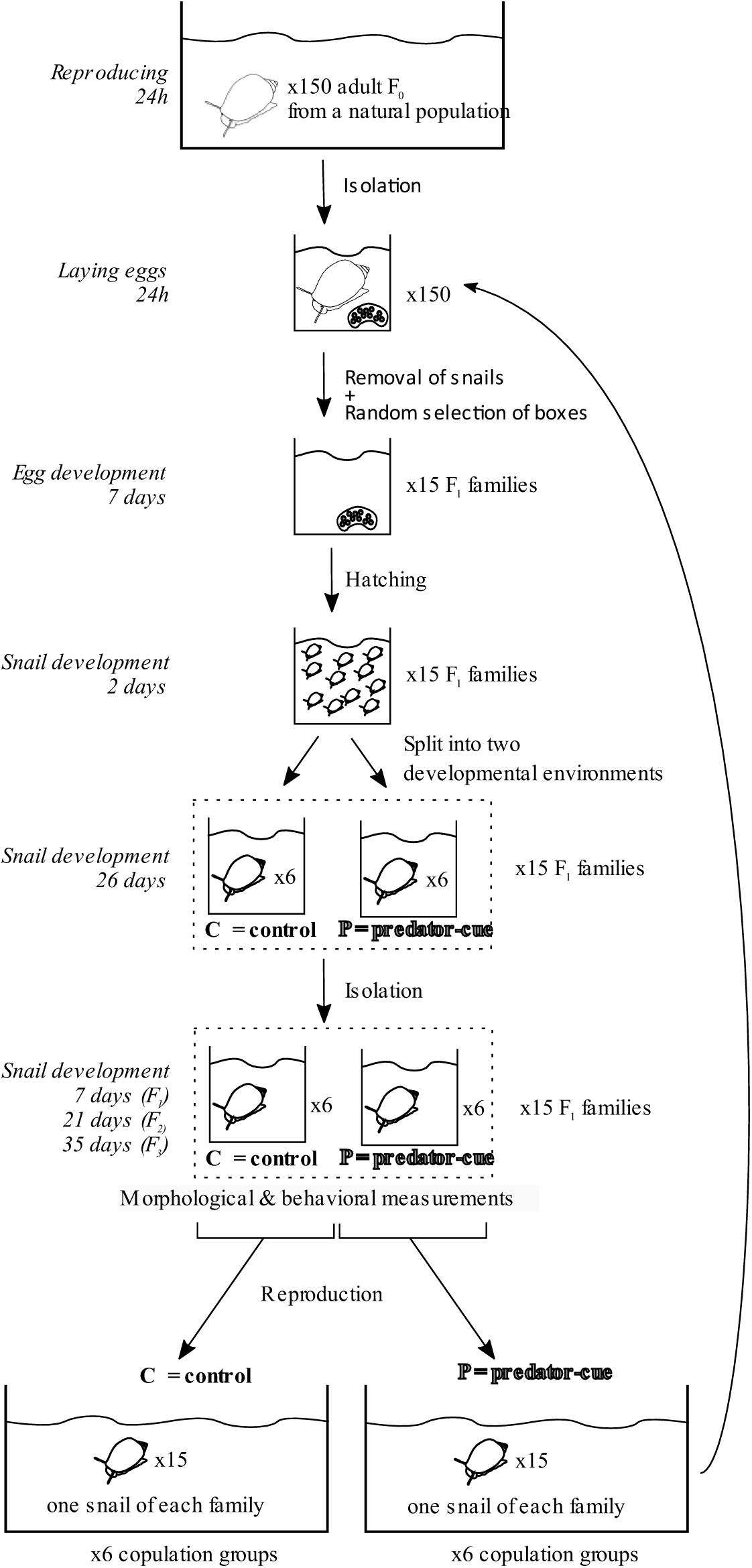

### Appendix 2. Principal component analysis for shell morphology

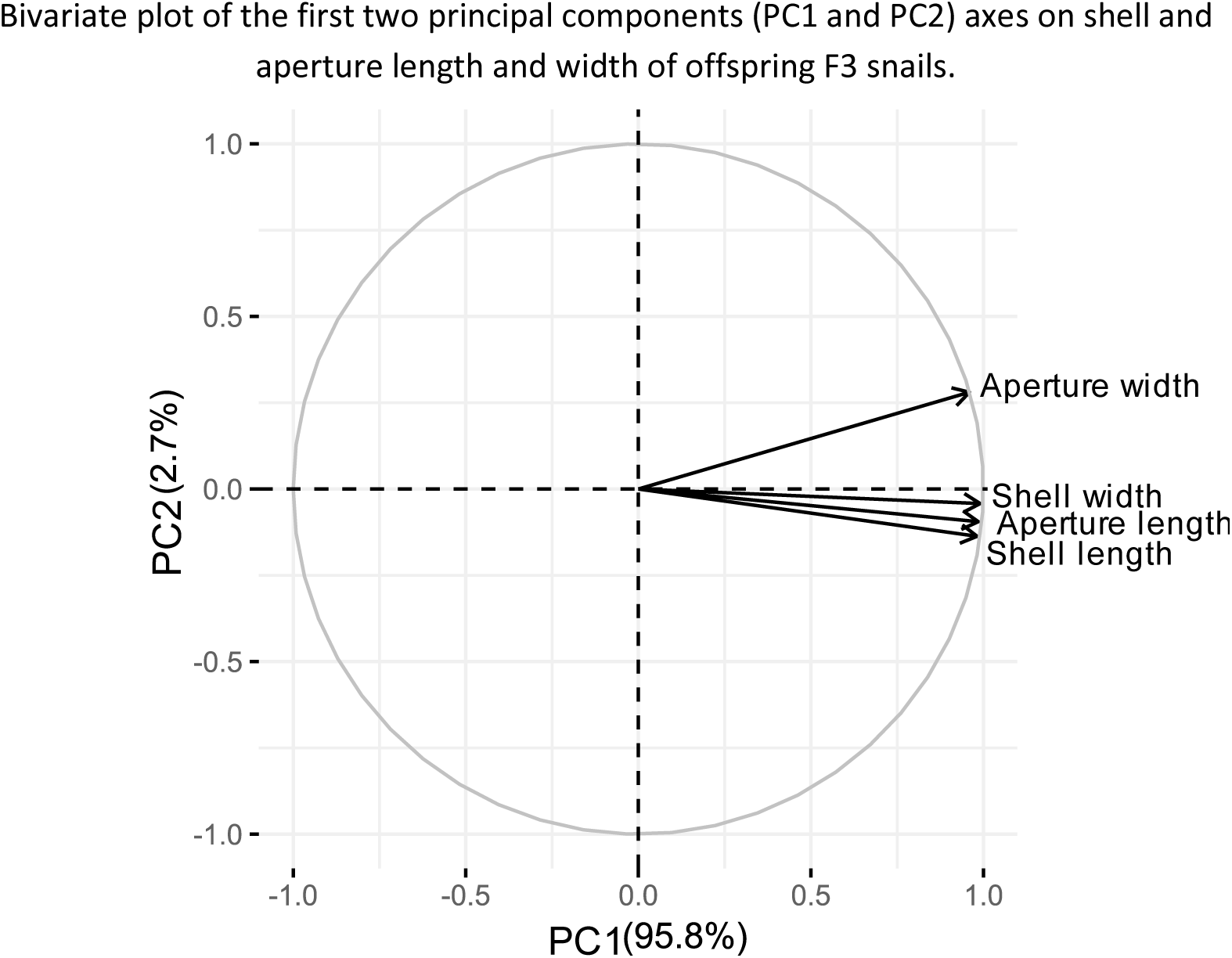

