## Supplementary Figure 2. Results PCA for shell morphology for "Transgenerational plasticity of inducible defenses: combined effects of grand-parental, parental and current environments" (from another version of this preprint: version 2, 2019-06-18)

### Supplementary figure 2: principal component analysis

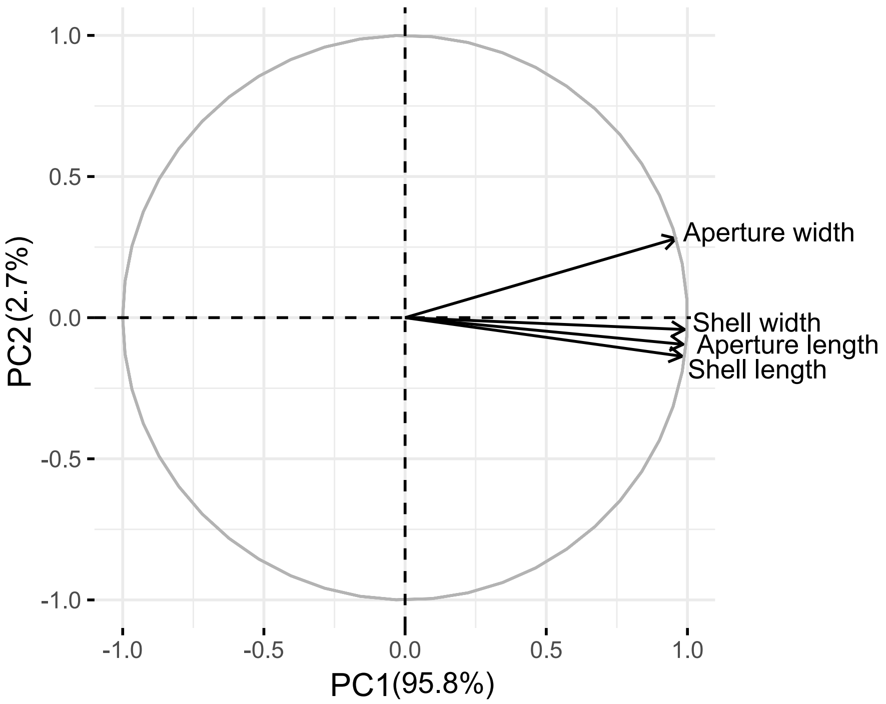


Bivariate plot of the first two principal components (PC1 and PC2) axes on shell and aperture length and width of offspring F3 snails.
